# Balanced mitochondrial function at low temperature is linked to cold adaptation in *Drosophila* species

**DOI:** 10.1101/2022.12.22.521616

**Authors:** Lisa Bjerregaard Jørgensen, Andrea Milena Hansen, Quentin Willot, Johannes Overgaard

## Abstract

The ability of ectothermic animals to live in different thermal environments is closely associated with their capacity to maintain physiological homeostasis across diurnal and seasonal temperature fluctuations. For chill-susceptible insects, such as *Drosophila*, cold tolerance is tightly linked to ion and water homeostasis obtained through a regulated balance of active and passive transport. Active transport at low temperature requires a constant delivery of ATP and we therefore hypothesize that cold-adapted *Drosophila* are characterized by superior mitochondrial capacity at low temperature relative cold-sensitive species. To address this, we investigated how experimental temperatures 19-1 °C affected mitochondrial substrate oxidation in flight muscle of seven tropical and temperate *Drosophila* species that represent a broad spectrum of cold tolerance. Mitochondrial oxygen consumption rates measured using a substrate-uncoupler-inhibitor-titration protocol showed that cooling generally reduced oxygen consumption of all steps of the electron transport system across species. Complex I is the primary consumer of oxygen at benign temperatures, but low temperature decreases complex I respiration to a much greater extent in cold-sensitive species than in cold-adapted species. Accordingly, cold-induced reduction of complex I correlates strongly with CT_min_ (the temperature inducing cold coma). The relative contribution of alternative substrates, proline, succinate and glycerol-3-phosphate increased as temperature decreased, particularly in the cold-sensitive species. At present it is unclear whether the oxidation of alternative substrates can be used to offset the effects of the temperature-sensitive complex I, and the potential functional consequences of such a substrate switch are discussed.

**Summary statement:** Mitochondrial oxygen consumption decreases at low temperature, particularly in cold-sensitive *Drosophila* species, which turn to oxidation of alternative substrates as complex I-supported respiration is impaired.

## Introduction

Most insects have limited capacity for physiological heat production and their body temperature is therefore primarily determined by the surrounding environmental conditions (Woods et al., 2015). As all biochemical and physiological processes are temperature-dependent it follows that environmental temperature will exert a strong influence on the physiological rates of insects, their performance and ultimately survival should temperatures become sufficiently extreme (Cossins and Bowler, 1987; Harrison et al., 2012; Hochachka and Somero, 2002). Given the overarching influence of temperature on physiological processes it is not surprising that interspecific variation in thermal tolerance is a strong determinant of biogeographical distribution of insects (Addo-Bediako et al., 2000; Bishop et al., 2017; Sunday et al., 2019). Limits for cold tolerance have been found to correlate particularly well with distribution in insects including the genus *Drosophila* that represents more than 1,400 “chill-susceptible” insect species distributed across tropical, temperate and subarctic climates (Kellermann et al., 2012; Kimura, 2004).

The cold biology of chill-susceptible insects is determined by their capacity to maintain physiological homeostasis when temperatures fluctuate seasonally, diurnally or during cold spells (MacMillan and Sinclair, 2011a; Nedved, 2000; Overgaard and Macmillan, 2017). Cold tolerance of chill-susceptible insects varies with acclimation (Weaving et al., 2022) and adaptation (Kellermann et al., 2012; Kimura, 2004) and it is well documented that chill tolerance is closely linked to the insect’s ability to maintain ion and water balance of their extracellular fluids (Koštál et al., 2004; Overgaard and Macmillan, 2017; Overgaard et al., 2021). Accordingly, chill-susceptible insects enter a state of neuromuscular paralysis (chill coma, CT_min_) at a temperature where they are unable to defend extracellular K^+^ balance within the CNS (Andersen et al., 2018; Armstrong et al., 2012), and chronic cold injury occurs when low temperature impairs osmoregulatory capacity to a degree that causes hemolymph hyperkalemia (Koštál et al., 2006; MacMillan and Sinclair, 2011b; Overgaard et al., 2021).

Because chill tolerance relies on the ability of insects to balance active and passive ion transport, it follows that the ability to fuel active transport at low temperature could represent an important cold adaptation. The critical thermal minima (CT_min_) is seemingly unaffected by oxygen availability at low temperature (Boardman et al., 2016; Stevens et al., 2010) and cold exposure, even below CT_min_, is usually not associated with a decrease in total ATP concentration (Colinet, 2011; Koštál et al., 2004; MacMillan et al., 2012; Williams et al., 2018). Nevertheless, it has been discussed whether mitochondrial capacity including the electron transport system (ETS) and associated oxidative phosphorylation is compromised/modified by low temperature in insects (Colinet et al., 2017; Lubawy et al., 2022; Menail et al., 2022; Wood and Nordin, 1980).

Exposure of *Drosophila melanogaster* mitochondria to 4 °C acutely decreases oxygen consumption rate relative to that measured at 25 °C in both control and cold-acclimated flies and chronic exposure to 4 °C further decreased the capacity for mitochondrial oxygen consumption and ATP synthesis (Colinet et al., 2017). However, the mitochondrial ATP/O ratio (ATP produced per molecule oxygen consumed) and respiratory control ratio (RCR – oxygen consumption associated with oxidative phosphorylation vs. proton leak) were only moderately affected by acute and chronic cold. These findings suggest that the effects of cold stress on mitochondrial capacity were caused by mitochondrial breakdown rather than altered function of the isolated mitochondria (Colinet et al., 2017). Nevertheless, the same study found a tendency for cold-acclimated *D. melanogaster* to defend ATP/O and RCR better during chronic cold. Similar tendencies were reported for cold-acclimated mayfly larvae that retained relatively higher respiration capacity and mitochondrial coupling (RCR) at low temperature compared to their warm-acclimated conspecifics (Havird et al., 2020). Cold-induced changes in insect mitochondrial capacity and coupling could also be associated with unbalanced temperature effects on different complexes in the electron transport system (Havird et al., 2020; Menail et al., 2022). For example, Menail et al. (2022) observed significant decreases in oxygen consumption rate (OCR) of complex I (CI-OXPHOS) in *D. melanogaster* and honeybees at low temperature (6-12 °C) compared to OCRs at a benign temperature (18 °C). The reduced OCR was partially compensated when alternative substrates (proline, succinate and glycerol-3-phosphate (G3P)) were added, suggesting that low temperature acutely alters the relative importance of ETS complexes (Menail et al., 2022). The findings discussed above suggest that insect mitochondrial function is modified or challenged by low temperature, but at present there is very limited knowledge of how cold adaptation is manifested at the functional level in mitochondria.

To investigate whether cold adaptation modifies mitochondrial capacity and function at low temperature, we examined the association between cold adaptation and mitochondrial respiration capacity in seven *Drosophila* species. The seven species of temperate and tropical *Drosophila* were chosen broadly from the phylogeny to represent independent examples of cold-sensitive and cold-tolerant species as attested by their marked difference in chill tolerance (CT_min_ ranging from 7.02 to - 1.94 °C). For each species mitochondrial function was studied at five temperatures (19-10-7-4-1 °C) using a substrate-uncoupler-inhibitor titration (SUIT) protocol to evaluate the contribution of individual mitochondrial complexes to the overall oxygen consumption rate. Given the temperature effect on biochemical processes and the requirement for sustained energy production to maintain homeostasis at low temperature, we hypothesized that 1) mitochondrial oxygen consumption will decrease with temperature, 2) the temperature effect on mitochondrial oxygen consumption and coupling will correlate with species cold tolerance, i.e., cold-adapted species will maintain higher OCRs and remain well-coupled at low temperature compared to cold-sensitive species, and 3) that the relative contribution of ETS complexes will be stable in cold-adapted species but may change with temperature in cold-sensitive species.

## Materials and methods

### Experimental animals

Mitochondrial respiration was measured in seven *Drosophila* species that were selected broadly from the phylogeny to include both cold-sensitive and cold-tolerant species from each of the two main subgenera *(Sophophora* (S) and *Drosophila* (D)). The species used are here listed in relation to their increasing level of cold tolerance (CT_min_ ± s.e.m.; subgenera, see below): *D. sulfurigaster* Duda 1923 (7.02 ± 0.11 °C; D), *D. bunnanda* Schiffer and McEvey 2006 (6.59 ± 0.07 °C; S), *D. teissieri* Tsacas 1971 (6.41 ± 0.16 °C; S), *D. melanogaster* Meigen 1830 (3.47 ± 0.12 °C, S), *D. obscura* Fallén 1823 (-0.13 ± 0.13 °C;S), *D. persimilis* Dobzhansky and Epling 1944 (−0.81 ± 0.12 °C; S) and *D. montana* Patterson and Wheeler 1942 (−1.94 ± 0.12 °C; D).

Flies were kept under common conditions (19 °C, 22:2 h light:dark) in 250-mL plastic bottles containing 35 mL oat-based Leeds medium (Andersen et al., 2015). Parental flies were transferred to new bottles twice a week to prevent high larval density, and eclosed flies for experiments were transferred to new bottles every 2-3 days. Only female flies of the age 3-12 d were used for measurements.

### Measurements of the critical thermal minimum CT_min_

Individual flies were placed in 5-mL glass vials, mounted to a rack and submerged in a tank connected to a cooling bath with ethylene glycol set to 19 °C (LAUDA-Brinkmann, NJ, USA). Temperature was gradually decreased by 0.1 °C min^-1^, and flies were checked for movement with increasing frequency as the flies started to show impaired motor function at low temperature. Once spontaneous movement stopped, vials were prodded, and flies were again checked for movement; the temperature where flies no longer responded was taken as the critical thermal minimum CT_min_. Measurements were made for n = 10 for each species except *D. obscura, D. melanogaster* and *D. teissieri* (n = 8, 9 and 9, respectively).

### High-resolution mitochondrial respirometry

Mitochondrial oxygen consumption rates were measured in permeabilized thoraces using the Oxygraph-O2K system (Oroboros Instruments, Innsbruck, Austria) and the associated software (DatLab v7.0.4.1, Oroboros Instruments). Measurements were performed at five temperatures: 19 °C (rearing temperature) and lower temperatures 10, 7, 4 and 1 °C to cover critical temperatures for most species. The protocol was similar to that used in Jørgensen et al. (2021), but is briefly outlined below.

### Permeabilization of thoraces

Preparation of thoraces was performed over ice. Flies were placed on ice to incapacitate them and with a scalpel and a pair of tweezers, the thorax was separated from the head, abdomen, wings and legs and then placed in ice-cold biological preservation solution [BIOPS; 2.77 mmol L^-1^ CaK_2_EGTA, 7.23 mmol L^-1^ K_2_EGTA, 5.77 mmol L^-1^ Na_2_ATP, 6.56 mmol L^-1^ MgCl_2_, 20 mmol L^-1^ taurine, 15 mmol L^-1^ Na_2_-phosphocreatine, 20 mmol L^-1^ imidazole, 0.5 mmol L^-1^ dithiothreitol and 50 mmol L^-1^ K-MES, pH 7.1 (Simard et al., 2018)]. Thoraces were gently punctured with a pair of fine-tipped forceps for initial mechanical permeabilization and then placed on a plate rotator (100 rpm) for 15 min in Eppendorf tubes with saponin-supplemented BIOPS (62.5 μg mL^-1^, prepared daily) for chemical permeabilization. To stop the chemical permeabilization, thoraces were transferred to Eppendorf tubes with ice-cold respiration medium [RESPI; 120 mmol L^-1^ KCl, 5 mmol L^-1^ KH_2_PO_4_, 3 mmol L^-1^ HEPES, 1 mmol L^-1^MgCl_2_ and 1 mmol L^-1^ EGTA, adjusted to pH 7.2 before adding 0.2% (w/v) fatty acid free BSA (Simard et al., 2018)] and placed on the plate rotator (100 rpm) for 10 min. Thoraces were then gently blotted dry on a tissue and weighed [MSE6.6S-000-DM micro balance (0.001 mg), Sartorius, Göttingen, Germany] before being placed in a small droplet of RESPI on parafilm over ice. The number of thoraces for each chamber was chosen considering the species thorax size and the experimental temperature (balancing a clear oxygen consumption signal with the risk of depleting chamber oxygen during the protocol). Generally, more thoraces were used for the smaller species (*D. melanogaster*, *D. teissieri* and *D. bunnanda*), and overall, the median mass used was 1.27 mg (1^st^-3^rd^ quartile: 0.98-1.80).

### Measurement of oxygen consumption rates

To calibrate the Oxygraph, temperature was set and 2.5 mL RESPI was added to each chamber after which chambers were closed with stoppers and excess RESPI was aspirated to ensure a chamber volume of 2 mL. Then, stoppers were lifted with the associated spacer to equilibrate the oxygen concentration to that of air, and once the OCR was stable at ± 1 pmol O2 s^-1^ mL^-1^, the chambers were calibrated with respect to the pertaining barometric and water vapor pressure (DatLab v7.0.4.1, Oroboros Instruments). Oxygen consumption rates were measured using a substrate-uncoupler-inhibitor-titration (SUIT) protocol (Jørgensen et al., 2021; Simard et al., 2018). To start the measurements, pyruvate (10 mmol L^-1^), malate (2 mmol L^-1^) and fly thoraces were added to each chamber to stimulate electron transport and proton pumping through complex I without coupling to oxidative phosphorylation (CI-LEAK). As insect flight muscle can have very high rates of oxygen consumption, chambers were supplemented with oxygen to increase the oxygen concentration to ≈ 650 μmol L^-1^ before closing. The SUIT protocol was then initiated, and each substance was injected once the OCR was stable. First, ADP (5 mmol L^-1^) was added to couple proton pumping to the oxidative phosphorylation by ATP synthase (CI-OXPHOS). Next, cytochrome *c* (10 μmol L^-1^) was injected to examine the integrity of the outer mitochondrial membrane (Kuznetsov et al., 2008), and in subsequent analysis, measurements with cytochrome *c*-induced increases in OCR exceeding 15 % were discarded. Then, proline (5 mmol L^-1^) was added to stimulate proline dehydrogenase (ProDH), a complex that transfers electrons to the Q-junction then complex III (CI+ProDH-OXPHOS), followed by succinate (20 mmol L^-1^) to stimulate electron transport through complex II (CI+ProDH+CII-OXPHOS). Then, *sn*-glycerol-3-phosphate (15 mmol L^-1^, G3P) was injected to stimulate electron transport through the mitochondrial G3P dehydrogenase (mtG3PDH), another complex transferring electrons to the Q-junction (CI+ProDH+CII+mtG3PDH-OXPHOS).

The uncoupler carbonyl cyanide 4-(trifluoromethoxy)phenylhydrazone (FCCP) was sequentially injected in doses of 1-2 μmol L^-1^ to examine whether the ETS could increase electron transport capacity when the proton gradient was uncoupled from oxidative phosphorylation. Injections of FCCP were made until the OCR no longer increased (FCCP-ETS). At this point, the main ETS complexes were sequentially inhibited; complex I by rotenone (0.5 μmol L^-1^), complex II by malonate (5 mmol L^-1^, prepared daily) and complex III by antimycin A (2.5 μmol L^-1^). The residual OCR remaining after complex inhibition originates from non-mitochondrial oxidative side reactions and was therefore subtracted from all measured OCRs before data processing. If oxygen concentration decreased below 180 μmol L^-1^ after inhibition, the oxygen concentration was increased before the next steps (same procedure as above). Following stabilization of the OCR, ascorbate (2 mmol L^-1^) and N,N,N’,N,-tetramethyl-p-phenylenediamine (TMPD, 0.5 mmol L^-1^) was injected to stimulate electron transport through cytochrome *c* which delivers electrons to CIV. After the OCR had peaked, sodium azide (20 mmol L^-1^) was injected to inhibit complex IV, and the system was left for 15-20 minutes with the OCR gradually decreasing to subsequently account for auto-oxidation of TMPD when calculating the maximal OCR of complex IV.

### Analysis of respiration data

Unless otherwise stated, OCRs are reported as mean±s.e.m. of mass-specific rates (pmol O2 s^-1^ mg^-1^ permeabilized thorax).

Three parameters were calculated from the measured OCRs. The OXPHOS-coupling efficiency (*j*_≈*p*_) was calculated as

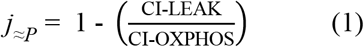

If OCR increased considerably after injection of ADP, *j*_≈*p*_ approaches 1, indicating a strong coupling between electron transport through complex I and oxidative phosphorylation. Contrarily, *j*_≈*p*_ approaching 0 indicates a weak coupling between these two ETS components.

The substrate contribution ratio (SCR) indicates the relative increase in OCR when proline, succinate or G3P was added and was calculated as:

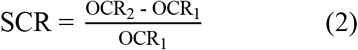

Here, OCR_1_ and OCR_2_ are the oxygen consumption rates before and after addition of the new substrate, respectively. If SCR = 0 there was no change in OCR, while SCR = 0.5 represents a 50 % increase, and SCR = 1 represents a doubling of the OCR following substrate injection. Note that this ratio is sensitive to the effect of the added substrate, but also to the level of OCR before addition of the substrate.

The effect of uncoupling (uncoupling control ratio, UCR) was assessed as:

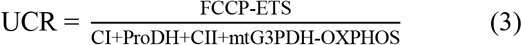

Here, FCCP-ETS and CI+ProDH+CII+mtG3PDH-OXPHOS are the maximal oxygen consumption rates when electron transport is uncoupled from and coupled to oxidative phosphorylation, respectively. When UCR = 1, the ETS is already at its maximal capacity for electron transport, while UCR > 1 indicates that uncoupling the ETS from ATP synthase (which utilizes the produced proton gradient) increases electron transport and the ETS was therefore limited by the downstream capacity of ATP synthase.

Species-temperature interactions on OCR was examined for each step of the SUIT protocol. To aid comparison of temperature responses in OCRs between species that can have different basal levels of oxygen consumption, the OCRs were standardized by dividing each OCR by the species-specific mean OCR of CIV at 19°C for a relative value (unitless). This reference value was chosen since, for most of our measurements, the ETS exhibits the highest OCR when complexes I-III are inhibited and CIV is stimulated, and moreover, 19°C had the highest OCR on average. Additionally, it is frequently observed that CIV has excess capacity compared to the ETS (Gnaiger et al., 1998; Pichaud et al., 2011).

To examine the correlation between mitochondrial function and species cold tolerance, OCRs of CI-OXPHOS and CI+ProDH+CII+mtG3PDH-OXPHOS were normalized to the mean of the specific rate at 10 °C. A linear regression on the OCRs at the two temperatures bracketing a 50 % decreased OCR relative to 10 °C gave proxies for temperature sensitivity of CI-OXPHOS and CI+ProDH+CII+mtG3PDH-OXPHOS, which were then analyzed in relation to species cold tolerance (CT_min_) using linear regressions. The thermal sensitivity quotient Q_10_ was calculated for CI-OXPHOS and CI+ProDH+CII+mtG3PDH-OXPHOS in the temperature interval 1-10 °C using the formula:

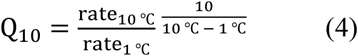

Where rate_10 °c_ and rate_1 °c_ is the respective mean rate of CI-OXPHOS or CI+ProDH+CII+mtG3PDH-OXPHOS at 10 and 1 °C, respectively.

### Statistics

Statistical data analyses were performed in R v4.2.1 (R Core Team, 2022).

Oxygen consumption was compared; i) between temperatures for each step of the SUIT protocol within species (mass-specific OCRs), ii) between species within temperature and substrate (standardized OCRs), and iii) as substrate contribution ratios between species within temperature. Furthermore, the OXPHOS coupling efficiency (*j*_≈*p*_) and the uncoupling control ratio (UCR) were compared between temperatures within species using one-way ANOVAs followed by a post hoc HSD Tukey’s test.

## Results

### Low temperature decreases oxygen consumption rate in a species-specific manner

To investigate how low temperature affects mitochondrial function, mass-specific OCR was measured at different steps of the ETS at five temperatures (1, 4, 7, 10 and 19 °C) in seven *Drosophila* species.

When oxygen consumption rates were compared between temperatures within species it was clear that lowering of temperature decreased OCR for all the examined steps of the ETS. This effect was significant across most of the species and experimental conditions, however, for some species (*D. persimilis, D. melanogaster* and *D. sulfurigaster*), CI-LEAK did not decrease significantly with decreasing temperature (Fig. 1, Table S1). The uncoupling ratio (UCR) based on FCCP-ETS/CI+ProDH+CII+mtG3PDH-OXPHOS was always above 1 irrespective of species and temperature, indicating that the electron transport was limited by the phosphorylation system, i.e., there was an untapped capacity for electron transport in the system (Table 1). In five of the seven species we found a slight tendency for UCR to increase with decreasing temperature, but this was only significant for *D. persimilis* and *D. melanogaster*. For one species, *D. teissieri*, UCR varied significantly with temperature (highest at 10 °C and lowest at 7 °C) (Table 1).

**Figure 1:**
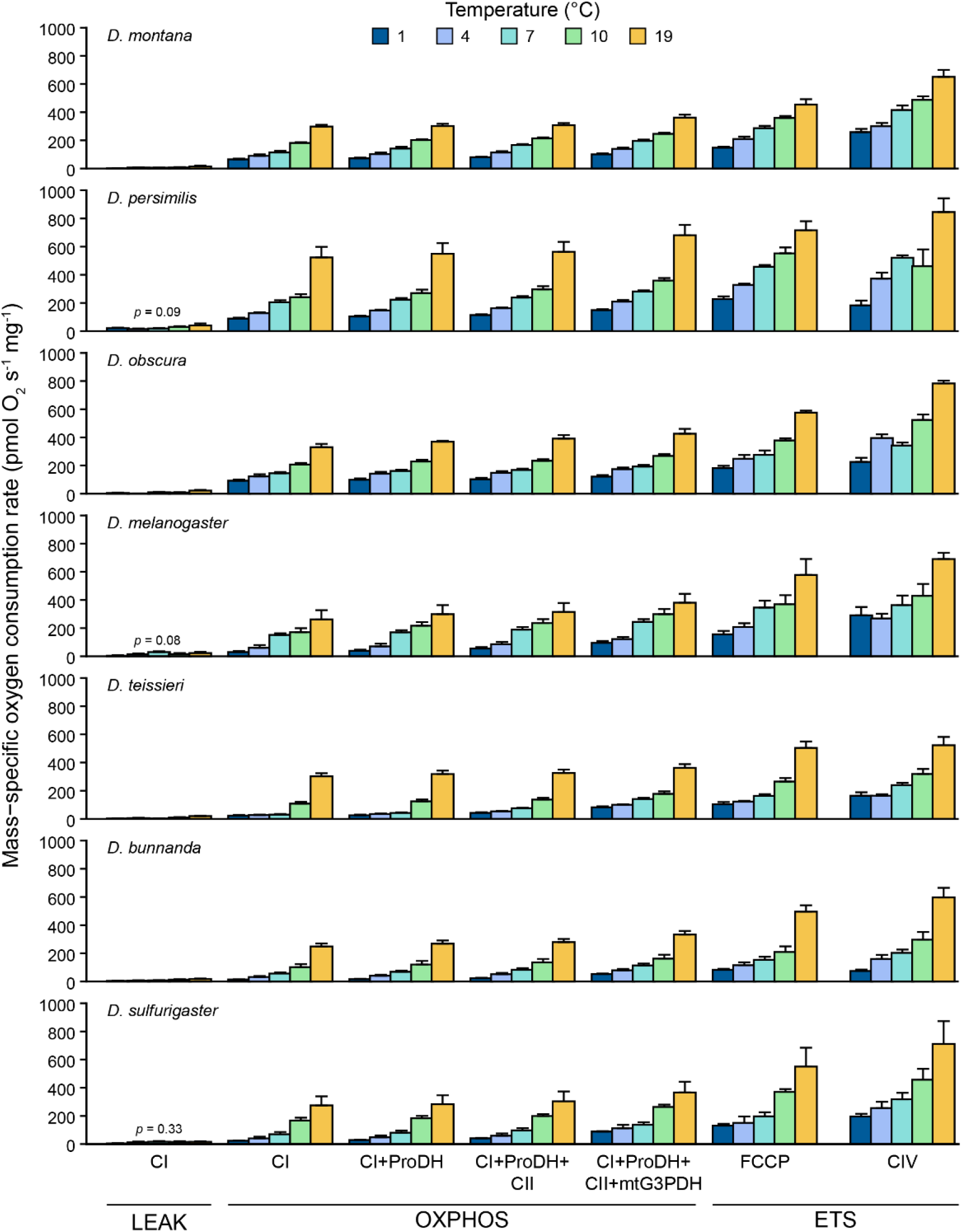
*Mass-specific oxygen consumption rate in permeabilized* Drosophila *thoraces at different temperatures*. *Oxygen consumption was measured in different states using a SUIT protocol; LEAK (without coupling to oxidative phosphorylation), OXPHOS (coupled to oxidative phosphorylation) and ETS (the electron transport system was uncoupled from oxidative phosphorylation). Oxygen consumption rates (OCRs) are reported as means±s.e.m., and for each step of the protocol (cluster of bars), OCRs were compared between experimental temperatures within species using a one-way ANOVA. All tests indicated significant effects of temperature (p < 0.05), except for CI-LEAK in* D. persimilis, D. melanogaster *and* D. sulfurigaster *(Table S1)*.

**Table 1:**
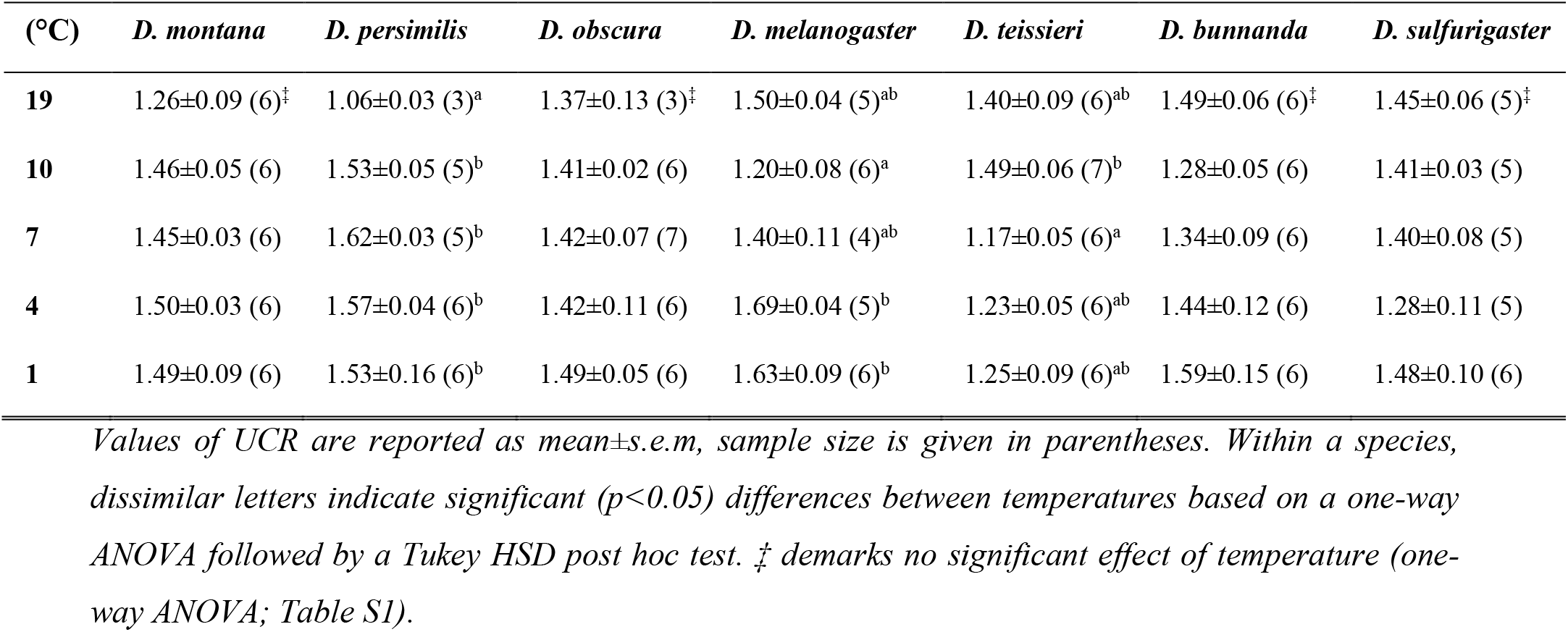
Uncoupling control ratio (UCR).

| (°C) | *D. montana* | *D. persimilis* | *D. obscura* | *D. melanogaster* | *D. teissieri* | *D. bunnanda* | *D. sulfurigaster* |
| --- | --- | --- | --- | --- | --- | --- | --- |
| 19 | 1.26±0.09 (6)<sup>‡</sup> | 1.06±0.03 (3)<sup>a</sup> | 1.37±0.13 (3)<sup>‡</sup> | 1.50±0.04 (5)<sup>ab</sup> | 1.40±0.09 (6)<sup>ab</sup> | 1.49±0.06 (6)<sup>‡</sup> | 1.45±0.06 (5)<sup>‡</sup> |
| 10 | 1.46±0.05 (6) | 1.53±0.05 (5)<sup>b</sup> | 1.41±0.02 (6) | 1.20±0.08 (6)<sup>a</sup> | 1.49±0.06 (7)<sup>b</sup> | 1.28±0.05 (6) | 1.41±0.03 (5) |
| 7 | 1.45±0.03 (6) | 1.62±0.03 (5)<sup>b</sup> | 1.42±0.07 (7) | 1.40±0.11 (4)<sup>ab</sup> | 1.17±0.05 (6)<sup>a</sup> | 1.34±0.09 (6) | 1.40±0.08 (5) |
| 4 | 1.50±0.03 (6) | 1.57±0.04 (6)<sup>b</sup> | 1.42±0.11 (6) | 1.69±0.04 (5)<sup>b</sup> | 1.23±0.05 (6)<sup>ab</sup> | 1.44±0.12 (6) | 1.28±0.11 (5) |
| 1 | 1.49±0.09 (6) | 1.53±0.16 (6)<sup>b</sup> | 1.49±0.05 (6) | 1.63±0.09 (6)<sup>b</sup> | 1.25±0.09 (6)<sup>ab</sup> | 1.59±0.15 (6) | 1.48±0.10 (6) |

To identify putative interspecific patterns in the response to lowered temperature, the OCRs were standardized relative to the mean of the oxygen consumption rate of complex IV at 19 °C for each species (Figs. 2 and S1). For CI-OXPHOS, when mitochondria were stimulated with pyruvate, malate and ADP, the relative OCRs revealed a clear pattern of species-specific temperature effects. Low temperature led to a larger reduction of CI-OXPHOS in the three cold-sensitive species (*D. teissieri, D. bunnanda* and *D. sulfurigaster*) relative to the more cold-adapted species (*D. montana*, *D. persimilis* and *D. obscura*) while the cosmopolitan species (*D. melanogaster*) had an intermediate response (Fig. 2A; one-way ANOVA followed by Tukey’s HSD *post hoc* test, *F*-statistics in Table S2). The species differences in temperature response are significant at the three lowest temperatures (1, 4 and 7 °C) and the pattern is particularly clear at 4 °C where the level of CI-OXPHOS for *D. melanogaster* is intermediate to the three cold-sensitive and tolerant species, respectively.

**Figure 2:**
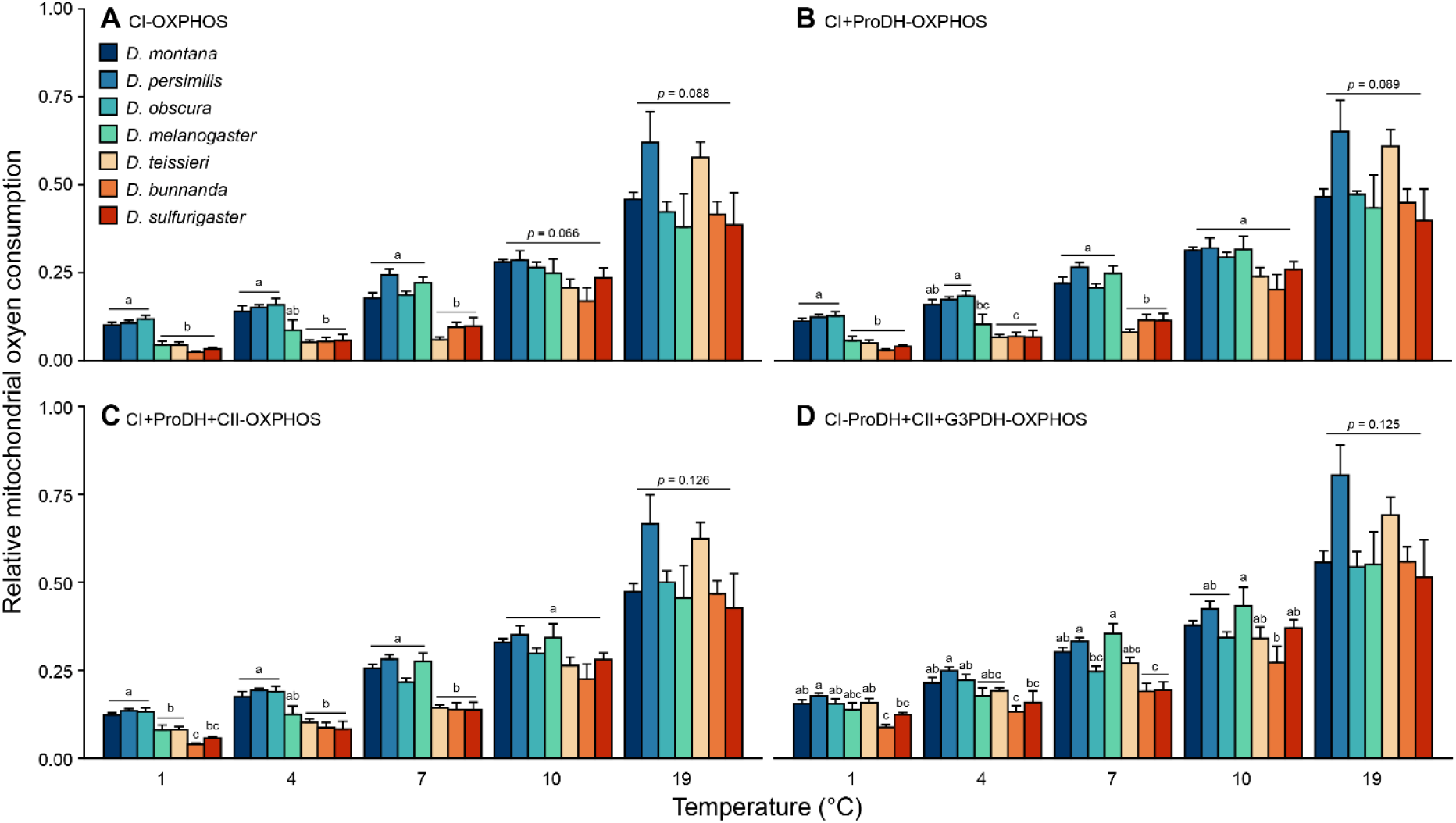
Relative oxygen consumption rates for comparison between species. Rates of A) CI-OXPHOS, B) CI+ProDH-OXPHOS, C) CI+ProDH+CII-OXPHOS and D) CI+ProDH+CII+mtG3PDH-OXPHOS were standardized to the average complex IV oxygen consumption rate at 19 °C. Within a temperature, dissimilar letters indicate significant (p < 0.05) differences between species based on a one-way ANOVA followed by a Tukey HSD post hoc test. p-values are shown from comparisons (one-way ANOVAs) that did not reveal a significant main effect of species (Table S2). Rates are presented as mean±s.e.m.

The interspecific pattern of temperature sensitivity generally persisted after addition of proline to stimulate CI+ProDH-OXPHOS (Fig. 2B) and after injection of succinate to fuel complex II (CI+ProDH+CII-OXPHOS, Fig. 2C). In both cases, the cold-adapted species had significantly higher relative OCRs than *D. melanogaster* and the cold-sensitive species at 1 °C, while at 4 °C, the relative OCRs of cold-adapted species were significantly higher than the cold-sensitive with *D. melanogaster* intermediate to the sensitive and tolerant species. Furthermore, at 7 °C the cold-sensitive species had significantly lower relative CI+ProDH-OXPHOS and CI+ProDH+CII-OXPHOS than the four more tolerant species while no interspecific differences were found for relative OCR at 10 and 19 °C.

Addition of G3P to stimulate mtG3PDH (CI+ProDH+CII+mtG3PDH-OXPHOS) changed the interspecific pattern (Fig. 2D) such that differences in relative OCR became smaller and were not significantly different between most species. Accordingly, G3P stimulated OCR more in the sensitive species such that total OCR levels became more similar (see below).

### Decreased CI-OXPHOS correlates with species cold tolerance

To examine the relationship between mitochondrial function at low temperature and species cold tolerance, CI-OXPHOS and CI+ProDH+CII+mtG3PDH-OXPHOS rates measured at 1-10 °C were normalized relative to the respective mean rate at 10°C. Normalization to 10 °C was chosen as the examined species are still able to move and exhibit normal neuromuscular function at this temperature (Andersen et al., 2015; MacLean et al., 2019) and because we found no major interspecific differences at this low, but tolerable temperature. In this analysis we estimated the temperature where the normalized OCR was reduced to 50 % of the activity at 10 °C using interpolation of the two bracketing temperatures (Fig. 3). Here it is seen that all species eventually experience a 50 % reduction in CI-OXPHOS with lowered temperatures, but there is a marked tendency for the cold-sensitive species to experience this reduction at higher temperatures than the cold-tolerant species (Fig. 3A). A similar, but less obvious pattern is found for CI+ProDH+CII+mtG3PDH-OXPHOS, where the cold-adapted species generally display the 50 % reduction at lower temperatures than the other species, but here the cold-sensitive *D. teissieri* also require a very low temperature to reduce the OCR (Fig. 3B). Further, we calculated the thermal sensitivity (Q_10_) of CI-OXPHOS and CI+ProDH+CII+mtG3PDH-OXPHOS for the interval 1-10 °C and found that CI-OXPHOS of cold-sensitive species generally was more sensitive to temperature decrease (higher Q_10_) than cold-adapted species, but that this difference was less clear for the maximal coupled oxygen consumption rate, CI+ProDH+CII+mtG3PDH-OXPHOS (Fig. 3A,B).

**Figure 3:**
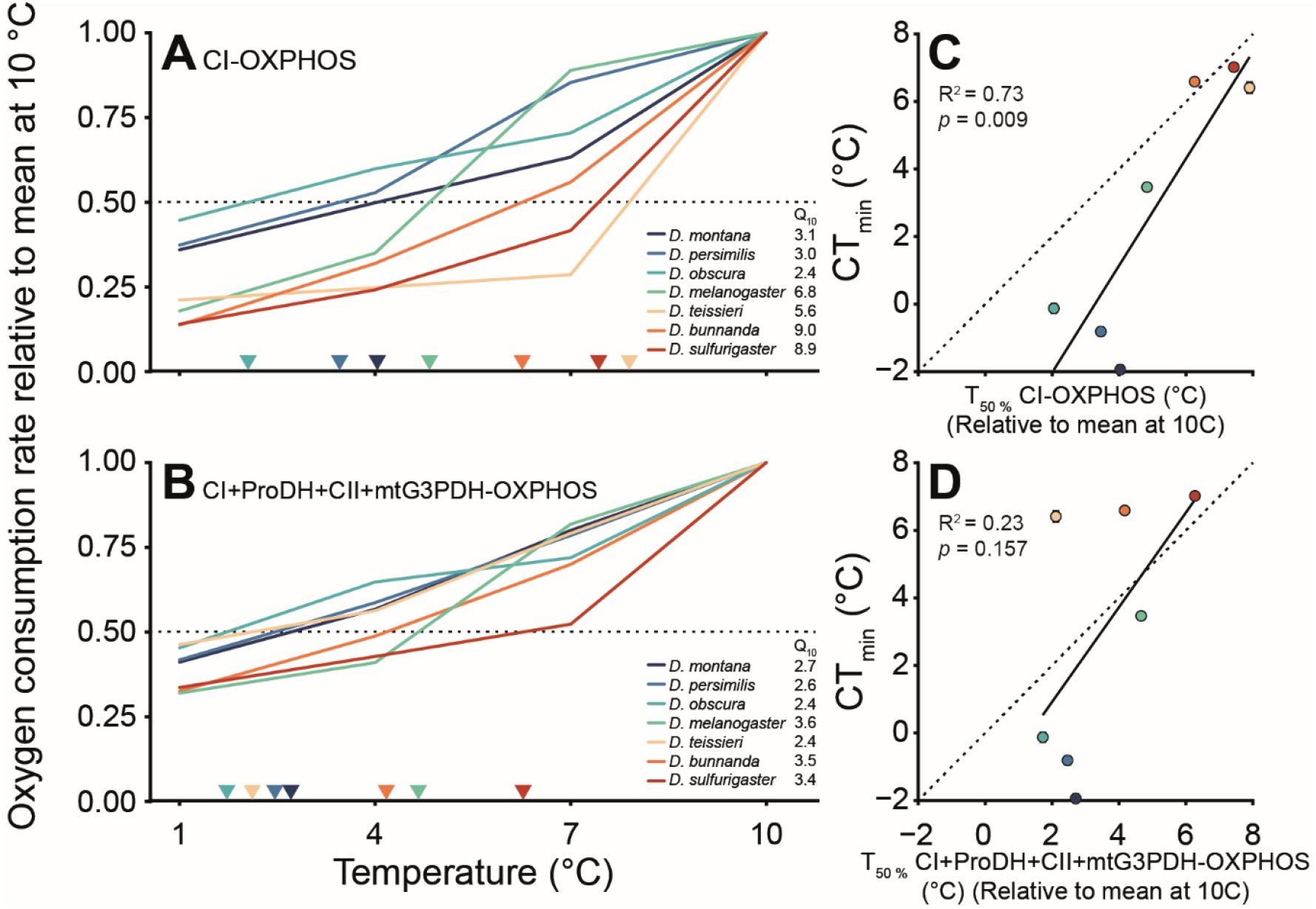
Temperature effects on relative oxygen consumption rates and correlation with species cold tolerance. Rates of A) CI-OXPHOS and B) CI+ProDH+CII+mtG3PDH-OXPHOS normalized to the respective mean OCR at 10 °C. The dotted line represents 50 % reduction in OCR relative to 10 °C. The temperature coefficient Q_10_ is calculated for the rates between 1-10 °C. Interpolated estimates of temperature where OCR is reduced by 50 % (T_50_ %) in C) CI-OXPHOS and D) CI+ProDH+CII+mtG3PDH-OXPHOS plotted against species mean CT_min_ along with the linear regression (solid line) and line of unity (dotted). Error bars (±s.e.m.) for the CT_min_ are hidden behind points.

To associate mitochondrial capacity to species cold tolerance, we regressed the index of mitochondrial function (the estimated temperature for 50 % reduction in normalized OCR for each species) to species cold tolerance, CT_min_. The resulting linear regression indicates that there is a strong correlation between the decrease in CI-OXPHOS and the level of cold tolerance (*R^2^* = 0.73; *F_1,5_* = 17.55, *p* = 0.009, Fig. 3C), while the correlation for the maximal coupled oxygen consumption CI+ProDH+CII+mtG3PDH-OXPHOS is not significant (*R^2^* = 0.23; *F_1,5_* = 2.77, *p* = 0.157, Fig. 3D).

Considering the significant association between CI-OXPHOS and cold tolerance we examined the temperature effect of coupling efficiency at the level of CI (*j*_≈*p*_), which describes the relation between CI-LEAK and CI-OXPHOS. Comparison of *j*_≈*p*_ between temperatures within species only revealed significant temperature effects in three of the seven species: *D. persimilis*, *D. melanogaster* and *D. bunnanda* (Table 2, Table S1). For *D. persimilis* and *D. bunnanda* the OXPHOS coupling efficiency observed at 1 °C was significantly lower than under control conditions at 19 °C and for *D. melanogaster*, *j*_≈*p*_ was significantly different between 4 and 19 °C, but increased at 1 °C.

**Table 2:** OXPHOS coupling efficiency (*j*_≈*p*_) at the level of complex I.

| (°C) | *D. montana* | *D. persimilis* | *D. obscura* | *D. melanogaster* | *D. teissieri* | *D. bunnanda* | *D. sulfurigaster* |
| --- | --- | --- | --- | --- | --- | --- | --- |
| 19 | 0.94±0.01‡ | 0.93±0.02<sup>a</sup> | 0.93±0.02‡ | 0.92±0.02<sup>a</sup> | 0.93±0.01‡ | 0.92±0.01<sup>a</sup> | 0.93±0.03‡ |
| 10 | 0.96±0.01 | 0.87±0.03<sup>ab</sup> | 0.96±0.01 | 0.92±0.03<sup>a</sup> | 0.89±0.02 | 0.85±0.04<sup>ab</sup> | 0.92±0.03 |
| 7 | 0.94±0.01 | 0.91±0.02<sup>a</sup> | 0.93±0.01 | 0.79±0.05<sup>ab</sup> | 0.89±0.03 | 0.86±0.02<sup>a</sup> | 0.77±0.04 |
| 4 | 0.92±0.02 | 0.88±0.03<sup>ab</sup> | 0.98±0.01 | 0.76±0.04<sup>b</sup> | 0.79±0.05 | 0.82±0.04<sup>ab</sup> | 0.69±0.13 |
| 1 | 0.95±0.02 | 0.77±0.04<sup>b</sup> | 0.94±0.02 | 0.85±0.04<sup>ab</sup> | 0.87±0.04 | 0.67±0.08<sup>b</sup> | 0.82±0.06 |

### Temperature-dependent shift in substrate oxidation varies with species

The observation that the fully stimulated maximal OXPHOS (CI+ProDH+CII+mtG3PDH-OXPHOS) had a weaker association with species cold tolerance than CI-OXPHOS suggests that addition of “alternative” substrates can partially compensate for the reduced CI-OXPHOS found particularly in the cold-sensitive species. To examine this explicitly we calculated substrate contribution ratios (SCRs) within temperatures for each species to investigate how much OCR increases with the addition of each added substrate. As the SCR reports the relative increase in OCR after injection of a substrate it is sensitive to both the increase in OCR, but also to the prevailing condition before addition of the substrate (i.e., if OCR is low before addition of the substrate, then SCR will increase more with a fixed increase in OCR as it reports the relative change).

Our analysis of SCR revealed significant temperature-dependent differences in mitochondrial substrate oxidation between species (Fig. 4, Table S2). Injection of proline generally had a low effect on the OCR, and there was no consistent pattern between species across temperatures (Fig. 4A). When succinate was injected at low temperatures (1-4 °C) there was a clear tendency for larger responses in OCR for the four most cold sensitive species compared to the three cold-tolerant species (Fig. 4B). Further, *D. teissieri* responded significantly more to the addition of succinate at 7 °C than all the other species. This suggests that succinate can partially compensate for the low CI-OXPHOS that characterizes the cold-sensitive species. A similar but even stronger increase in SCRs was found following addition of G3P (Fig. 4C) where cold-adapted species again responded relatively more to this substrate at 1 and 4 °C (also at 7 °C for *D. teissieri*). At 10 and 19 °C, SCRs were generally low across species for the three alternative substrates, again highlighting that the SCR is sensitive to the pertaining OCR before injection of alternative substrates (at 10 and 19 °C CI-OXPHOS is not compromised and accordingly the starting point for the calculation is high).

**Figure 4:**
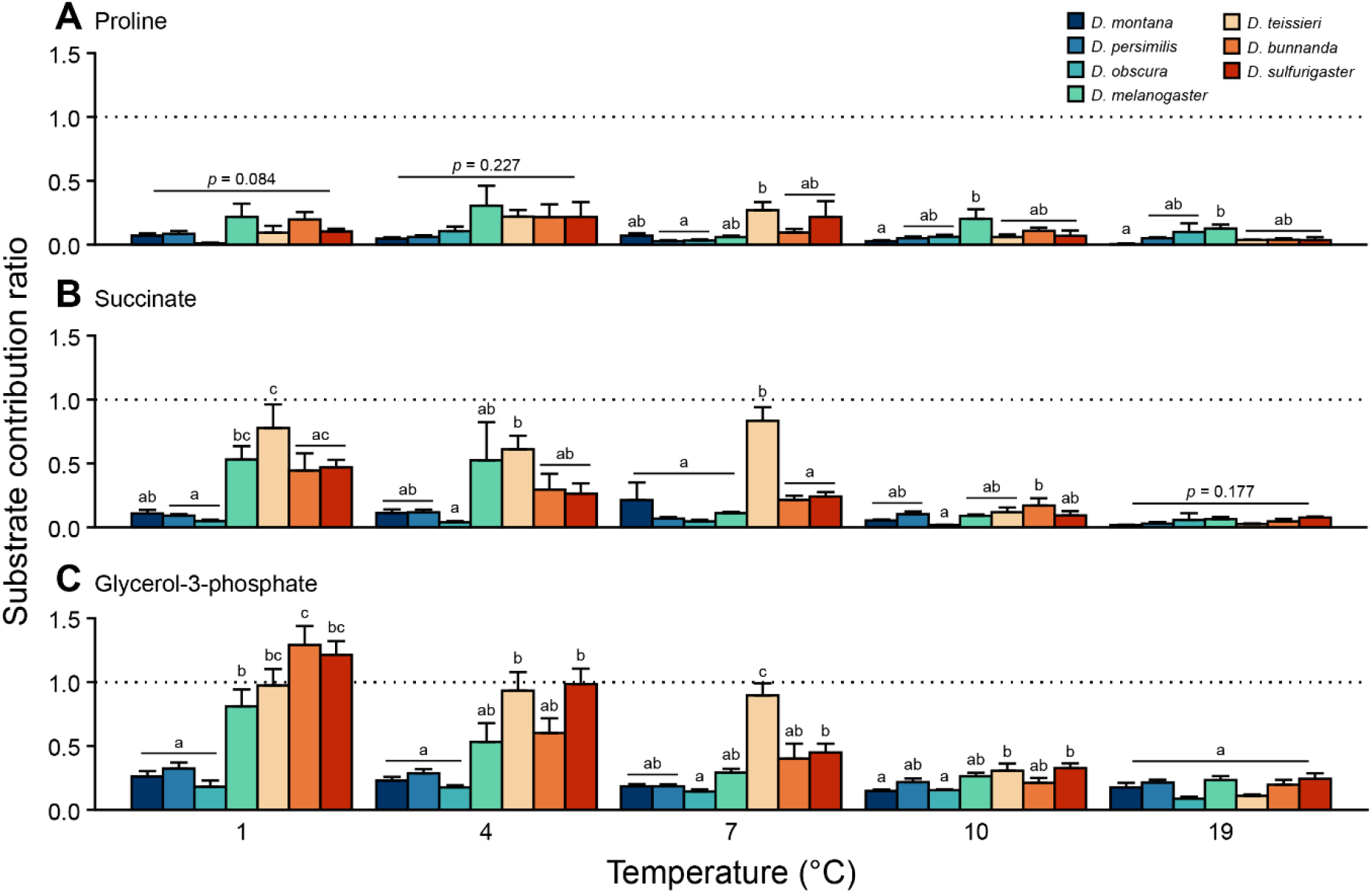
Substrate contribution ratios. Ratios (mean±s.e.m.) calculated for the increase in oxygen consumption following injection of A) proline, B) succinate and C) glycerol-3-phosphate. If SCR = 0, the substrate did not change the oxygen consumption rate, while SCR = 1 (dotted line) indicates a doubling in rate with substrate use. Within a temperature, dissimilar letters indicate significant (p < 0.05) differences between species based on a one-way ANOVA followed by a Tukey HSD post hoc test. p-values are shown for comparisons (one-way ANOVAs) that did not reveal a significant main effect of species (Table S3).

## Discussion

This study examined the acute effects of low temperature on mitochondrial function in permeabilized flight muscle of seven species of *Drosophila* that represent different levels of cold tolerance. Overall, our results show that mitochondrial oxygen consumption rate is strongly influenced by temperature; all seven species had reduced oxygen consumption at the lower temperatures (1-7 °C) compared to the higher temperatures (10-19°C) (Fig. 1). Notably, cold-sensitive species were characterized by larger reductions in complex I-supported respiration at low temperature than cold-adapted species (Figs. 2–3). As a result, these cold-sensitive species may rely more on oxidation of alternative metabolic substrates (Fig. 4).

### Temperature effects on complex I correlate with species cold tolerance

In accordance with expectations of normal temperature effects of biochemical processes (Cossins and Bowler, 1987; Hochachka and Somero, 2002; Jørgensen et al., 2022) we found that lowering the temperature decreased oxygen consumption rates of all complexes in the ETS. This temperature response was similar across all seven species when analyzed in the temperature interval between 10-19°C, a temperature range where all species are able to defend neuromuscular function. Thus, mitochondrial respiration rate decreased with an average Q_10_ of 2.1 and 2.9 for CI-OXPHOS and CI+ProDH+CII+mtG3PDH-OXPHOS, respectively, between 10-19 °C. Although the capacity for mitochondrial respiration is not directly comparable to resting metabolic rate at the organismal level (Davison, 1971; Tribe and Bowler, 1968), we note that the thermal sensitivity of mitochondrial respiration at these permissive temperatures is very similar to the average Q_10_ of 2.7 found for standard metabolic rate between 10-20 °C across more than 60 species of temperate and tropical *Drosophila* (Messamah et al., 2017).

The interspecific patterns of mitochondrial thermal sensitivity change markedly when oxygen consumption rates are measured in the range of temperatures from 10-1 °C. This range of temperatures will initially challenge the physiological performance of the three cold-sensitive species (*D. sulfurigaster*, *D. bunnanda* and *D. teissieri*) that enter cold coma at 6-7 °C and subsequently cold will stress the “intermediate” species (*D. melanogaster*) that enters cold coma at 3.5 °C. In contrast, the three tolerant species (*D. obscura*, *D. persimilis* and *D. montana*) only enter chill coma once temperatures have decreased below 0 °C. In the temperature interval between 10 and 1 °C we found clear differences in the dynamics and capacity of the mitochondrial respiration rate associated with species cold tolerance. This was particularly evident with respect to the reduction in complex I-supported respiration which was much more pronounced in the cold-sensitive species (Figs. 1–2). Here, Q_10_ of CI-OXPHOS was 7.6 (range: 5.6-9.0) for the four most cold sensitive species while the three cold-adapted species retained “normal” thermal sensitivity (Q_10_ = 2.9; range 2.4-3.1) (Fig. 3A). A similar change in thermal sensitivity of complex I-driven respiration was recently reported for *D. melanogaster* with Q_10_ ~ 10 at temperatures between 6-12 °C and Q_10_ ~ 1.9 between 12-18 °C (Menail et al., 2022). The association between cold tolerance and preservation of CI-OXPHOS is highlighted by the strong correlation found between the temperature estimated to induce a 50 % reduction of CI-OXPHOS relative to the rate at 10 °C (an index for decreased CI-OXPHOS capacity) and our measure of cold tolerance, CT_min_ (Fig. 3A,C). While this analysis suggests that preservation of CI-OXPHOS is important for cold tolerance it is important to emphasize that there is no direct mechanistic link to associate a particular rate of CI-OXPHOS (i.e., a specific rate in pmol s^-1^ mg^-1^) directly to the onset of CT_min_. Our findings simply suggest that thermal sensitivity of CI-OXPHOS is higher than that of organismal metabolic rate, which may cause unbalanced mitochondrial function as temperature decreases. CI-OXPHOS is the key contributor to the proton gradient across the inner mitochondrial membrane and the observed correlation suggests that the activity of this complex in the ETS may be of importance to support the capacity for ATP production. The few studies that have investigated ATP/O ratios at low temperature in insects have found these ratios to be well-defended at low temperature (Colinet et al., 2017; Lubawy et al., 2022; Wood and Nordin, 1980), but considering the differences in CI-OXPHOS found between species in the present study, it would be relevant to further examine the association between CI-OXPHOS and ATP/O ratio.

Interestingly, we recently found a similar strong association between failure of CI-OXPHOS and heat tolerance in six species of *Drosophila* (Jørgensen et al., 2021) again suggesting that preservation of CI-OXPHOS is important for homeostasis and thermal tolerance in *Drosophila*. It is likely, however, that the failure of CI-OXPHOS at high and low temperature is caused by different underlying mechanisms. At stressful high temperature the significant decrease in CI-OXPHOS is associated with a dramatic hyperthermic breakdown (Jørgensen et al., 2021). Such an acute disruption is not found at low temperature, and we instead suggest that the gradual decline in CI-OXPHOS at low temperature simply indicate that species are endowed with different thermal sensitivities that disproportionally reduces CI-OXPHOS in cold-sensitive compared to cold-tolerant species (Fig. 3A).

### Mitochondrial flexibility: Maintaining high oxygen consumption by oxidation of alternative substrates

Associated with the cold-induced suppression of CI-OXPHOS, we found that lowered temperature changed the relative contribution of mitochondrial substrate oxidation in all species *Drosophila*. The substrates succinate and G3P were of higher importance and had higher substrate contribution ratios (SCR) at low temperatures (1-7 °C) compared to high temperatures (10-19 °C) (Fig. 4B,C), while the effect of proline was generally low and uniform across temperatures (Fig. 4A). Importantly, the temperature effect on mitochondrial substrate oxidation differed significantly between species in relation to cold tolerance such that cold-sensitive species responded more to the addition of succinate and G3P at low temperatures. However, the SCRs are calculated from the preceding absolute OCR, and the low rates of CI+ProDH-OXPHOS and CI+ProDH+CII-OXPHOS prior to addition of succinate and G3P (as will be characteristic of cold sensitive species), will accordingly inflate the calculated SCRs for the same absolute increase. Thus, when examining the absolute increases in OCR after succinate and G3P at low temperatures (Fig. 1) we did not observe marked differences between species. Nevertheless, because the respiration rates of alternative substrates are less sensitive to lowered temperature across all species, we observed that the relative maximal OCR would generally offset the species differences in OCR that stem from species-effects on CI-OXPHOS (Figs. 2–3). A similar compensation with alternative substrates at low temperature has also been found with additions of succinate and G3P in *D. melanogaster* and honeybees (Menail et al., 2022), or with addition of succinate in warm-acclimated mayfly larvae (Havird et al., 2020) and a planarian (Scott et al., 2019). Thus, when considering Q_10_ for maximal OCR coupled to phosphorylation (after G3P injection) between 10 and 1°C we found more similar thermal sensitivities (Q_10_ for the four most sensitive species = 3.2 and Q_10_ for the three most tolerant species = 2.6) (Fig. 3B), which is also in line with Menail et al. (2022) who found Q_10_ ~ 2.3 between 6-12 °C in *D. melanogaster* when all substrates were available. Compensation using alternative substrates were also found following failure of CI-OXPHOS respiration at high temperature in six *Drosophila* species where particularly mitochondrial G3P oxidation served to compensate for a decreased CI-OXPHOS at high stressful temperatures (Jørgensen et al., 2021). At the genetic level, genes involved in the glycerophosphate shuttle (the mitochondrial shuttle that G3P participates in, which is essential for maintaining redox balance in insect flight muscle (Sacktor, 1975)) have been found to be more expressed at higher latitudes in *D. melanogaster* (Lavington et al., 2014), further suggesting that G3P oxidation may be important at low environmental temperatures. Finally, a study of the arctic blowfly and the temperate housefly found that G3P-supported respiration (with rotenone inhibiting complex I) was maintained at temperatures down to 2 °C (Wood and Nordin, 1980). Although the temperature sensitivity (activation enthalpy) increased markedly in both species below 11.5 °C the cold-tolerant arctic blowfly displayed a smaller increase in thermal sensitivity, again supporting the potential importance of G3P as an alternative substrate at low temperature (Wood and Nordin, 1980).

A switch in substrate oxidation associated with decreased function of complex I may alter the amount of reactive oxygen species (ROS) produced. The mitochondrial glycerol-3-phosphate dehydrogenase is a potent producer of superoxide, while complex I mainly contributes through reverse electron flow (Miwa et al., 2003). Reverse electron flow is possible if there is a high membrane potential and little substrate available for complex I, which is supported by the observation that mild uncoupling or inhibition of complex I by rotenone strongly reduces superoxide formation (Miwa et al., 2003). As complex I is a large contributor to the membrane potential, reducing its activity in the ETS may counter the adverse production of ROS, but on the other hand the increased reliance on G3P oxidation may increase the ROS formation from mtG3PDH, which additionally is less sensitive to uncoupling (Miwa et al., 2003). Future studies should therefor consider whether the reliance on G3P versus complex I substrates is associated with an increased ROS production which could contribute to the effects of cold stress.

### Temperature effects on coupling and implications for phosphorylation capacity

A high degree of OXPHOS coupling, which indicate the effect of coupling oxygen consumption to oxidative phosphorylation relative to that used to offset proton leak, has been associated with a high mitochondrial efficiency for ATP production, although the measure itself does not encompass ATP production efficiency directly (Colinet et al., 2017; Gnaiger, 2014; Salin et al., 2018). In a study on an arctic blowfly (Wood and Nordin, 1980), the ratio between mtG3PDH-OXPHOS and the mtG3PHD-LEAK, corresponding to the OXPHOS coupling efficiency reported in the present study, decreased markedly below 10 °C. Meanwhile, the efficiency of the phosphorylating system was relatively unaffected by temperature; it only decreased 10 % between 22-2 °C (Wood and Nordin, 1980). Unfortunately, the study by Wood and Nordin (1980) did not examine phosphorylation efficiency in the more cold sensitive housefly nor did they measure complex I respiration, making direct comparison to the present study difficult. In the present study we found that OXPHOS coupling efficiency was generally maintained at a high level, despite the decreased CI-OXPHOS at low temperature (Table 2). Although there was a tendency for a decreased coupling at lower temperature in some of the species, there was no correlation with cold tolerance. This suggests that proton leak does not pose a significant expense in terms of oxygen consumption at low temperature.

Uncoupling the ETS from phosphorylation increased the mitochondrial oxygen consumption (UCR > 1) across all temperatures and in all species. This indicates that the phosphorylating system is not able to utilize the full capacity of the electron transport system and therefore limit/control the flux through the ETS (Table 1). Thus, high UCRs indicate that ATP synthesis capacity (ATP synthase) or the capacity of the adenine nucleotide translocase (ANT), which switches ATP for ADP across the inner mitochondrial membrane, are limiting respiration rate (Pichaud et al., 2011). The UCRs found in the present study were generally well-above 1, and slightly higher than reported in other insect studies examining the effect of temperature on mitochondria (range: 0.85-1.5) (Chamberlin, 2004; Jørgensen et al., 2021; Menail et al., 2022). While there was a tendency for increased UCRs at lower temperatures, this was rarely supported statistically and seemingly not associated with species tolerance (Table 1).

### Conclusions

This study examined the effect of low temperature on mitochondrial function, and how interspecific differences in mitochondrial function is related to cold tolerance of *Drosophila* chosen broadly from the phylogeny. As hypothesized, mitochondrial respiration rates decreased with lower temperature in all species, but the dynamics of this response differs considerably between cold-adapted and cold-sensitive species (Fig. 1). Most notably, the reduction of complex I-supported respiration was larger and occurred at a higher temperature in cold-sensitive species (Figs. 2–3), and we therefore found a strong correlation between the maintenance of CI-OXPHOS at low temperature and the species cold tolerance. Cold-sensitive species were able to partially compensate their lowered complex I-supported respiration by increased reliance on alternative substrates, particularly G3P (Fig. 4). Together, our findings suggest that cold adaptation is manifested in a more stable and well-regulated reduction in mitochondrial metabolism, particularly complex I-supported respiration, while cold-sensitive species are forced to rely more on alternative metabolic substrates at low temperature which may potentially disrupt optimal mitochondrial function. Future studies should examine whether the temperature-induced change in substrate oxidation affects the species ROS production and/or their ability to maintain sufficient ATP production, to support physiological homeostasis, at low temperature.

## Acknowledgements

The authors would like to thank Kristoffer Neldeborg Jensen for help with Oroboros measurements and Elin Ellebæk Petersen for help with reagent preparation.

## Data availability

Data will be made available in a Zenodo repository upon acceptance.

